# A Deep Dive into the Cognitive Soundscape of Flow: Finding Your Groove

**DOI:** 10.64898/2026.05.13.724953

**Authors:** Benjamin Bartling

## Abstract

Flow state, characterized by optimal engagement and performance, represents a key concept in understanding human performance and cognitive resource allocation. Grounded in Csikszentmihalyi’s and Sherry’s flow theory and the Limited Capacity Model of Motivated Mediated Message Processing (LC4MP), this study investigated physiological and neural correlates of flow state during a simulated driving task under different music conditions and difficulty levels. Using a 2 × 3 factorial design with 20 participants, this study examined self-selected versus non-self-selected music across three difficulty levels, testing the relationship between task switching, cognitive resource allocation, and flow state. Physiological measures included heart rate and EEG (alpha/theta power) using a 4-channel Muse 2 headband, alongside a self-report measure of flow experience. Hierarchical linear modeling revealed significant physiological changes during self-selected music: heart rate decreased (*β* = −5.15, *p* < .001), while alpha (*β* = 5829.77, *p* < .001) and theta power (*β* = 7637.24, *p* < .001) increased. Task difficulty also showed significant effects, with heart rate decreasing during hard (*β* = −6.70, *p* < .001) and moderate (*β* = −3.40, *p* = .001) conditions. In particular, while physiological measures showed robust changes, the self-reported flow state did not reach significance. Task switching rates showed significant decreases during self-selected music (*β* = −0.86, *p* < .001) and hard difficulty (*β* = −0.61, *p* < .001), supporting the LC4MP framework’s predictions regarding cognitive resource allocation. These findings demonstrate how task switching and cognitive resource allocation relate to flow state induction. The results highlight the importance of multimodal measurement approaches and demonstrate that personal relevance through music selection and task difficulty significantly influence physiological and neural responses during performance. Future research should employ more comprehensive measurement approaches to better capture the complexity of flow-related neural activity and its relationship to task switching and cognitive resource allocation.

## 1. Introduction

Sound connects us. It is continuous, integrative, and expansive; it shapes who we are and how we connect to the world around us. Music’s profound impact transcends mere enjoyment. It holds the power to emotionally move humans to states of ecstasy where all sense of self, time, and space can be disregarded. It has permeated human experience across cultures and throughout history, weaving itself into the essence of our lives.

Most people have favorite musicians who possess an uncanny ability to move listeners to deep emotional levels. These individuals can invoke a state in which the listener loses all perception of self, time, and space. It is not the equipment utilized by such musicians, but rather their brain, body, and lived experiences that refine their unique sound over time. Thus, an artist’s tone and timbre serve as a medium through which they communicate, providing the potential to resonate with millions of individuals.

In every known culture, the ordering of sound in ways that please the auditory system has been used extensively to improve quality of life. One of the oldest and perhaps most popular functions of music is to focus listeners’ attention on patterns appropriate to a desired mood (Diaz, 2011). Music, as a form of organized auditory information, has the capacity to structure mental activity and reduce what Csikszentmihalyi, 1990 termed “psychic entropy,” the mental disorder that arises when irrelevant stimuli interfere with goal-directed thought. By occupying input channels with structured, emotionally engaging information, music can help mitigate boredom and anxiety while setting the stage for deeper cognitive engagement.

When individuals are attentively engaged, music can serve as a gateway to flow by organizing consciousness around a coherent sensory and emotional experience (Pates et al., 2003). This capacity is especially salient when individuals self-select music that is both complex and personally significant, as such choices may enhance analytical engagement and increase the propensity for entering a flow state. In this way, music listening becomes not merely a passive experience but an active, motivationally relevant process that aligns with both the appetitive system described in LC4MP and the conditions necessary for flow.

Music also offers multiple pathways through which well-being may be cultivated via intangible emotional expressions that activate reward systems in the brain and facilitate innovation and resonance (Kraus and Slater, 2015). However, a critical question remains: can music listening actively induce or contribute to the coveted flow state? Furthermore, does music-induced flow enable the channeling of emotions?

For the purposes of this project, the conceptualization of music does not focus on musical performers or musicians themselves, but rather on music listening as an applied aspect of music cognition in the broader population. To contextualize this perspective, the enactive approach to cognition proposed by Varela (1991) suggests that experience is co-created through reciprocal interactions among the mind, body, and environment. Within these interactions, the body operates through several closely related modalities. One such modality, termed “sensorimotor coupling,” allows individuals to perceive and attune themselves to the environment in ways that facilitate embodied interaction (Nijs et al., 2012).

Recent literature has neglected interindividual differences in musical listening experiences (Loepthien and Leipold, 2021). One study conducted by Loepthien and Leipold found a stronger relationship between flow experiences and subjective well-being among individuals with highly flexible self-concepts. Their findings demonstrated that flow was positively correlated with musical experience, providing evidence for the relationship between flow, musical practice, and musical listening experiences.

Prior research has examined flow in terms of situational and individual factors that facilitate its emergence, as well as its relationship with psychophysiological processes (Manzano et al., 2010; Pates et al., 2003; Sinnett et al., 2020; Valenzuela et al., 2018). However, most studies have focused on musical performance contexts rather than music listening, with only a limited number investigating individual listening experiences (Chirico et al., 2015; Pates et al., 2003).

Enhanced attentiveness has long been proposed as a defining characteristic of heightened affective experiences (Csikszentmihalyi, 1990; Krippner, 1972; Madsen and Geringer, 2008; Maslow, 1968). Maslow (1968) described such states as “peak experiences,” intense psychological moments characterized by feelings of profound happiness and fulfillment. Similarly, Csikszentmihalyi (1990) explained that intense enjoyment resulting from engagement in intrinsically rewarding tasks emerges partly from an individual’s degree of immersion in the activity itself. This experience is known as flow (Sherry, 2004).

Research suggests that music may provide a pathway toward unlocking flow states (Peifer and Engeser, 2021). Flow, a state coveted across disciplines, is characterized by complete absorption in an activity accompanied by heightened focus, intrinsic motivation, and optimal performance (Csikszentmihalyi et al., 2014). The present study investigates the intersection of music, cognition, task switching, and flow, specifically exploring how music interacts with the brain to facilitate the emergence of flow states.

Existing research clearly suggests that musical activities are conducive to flow, particularly when skills are appropriately matched with challenges, goals are clearly established, and contingent feedback is available (Nijs et al., 2012). However, limited attempts have been made to investigate heightened attention independently from other variables, and little effort has focused specifically on flow during music listening (Loepthien and Leipold, 2021). Accordingly, the present study examines relationships among flow, attention, and music listening using adaptations from research on cognition and attention.

Some scholars accept the notion that music is fundamentally embodied (Leman, 2007); however, the nature of flow remains comparatively underdeveloped theoretically. Sensorimotor coupling may provide a mechanism underlying the relationship between flow and music (Nijs et al., 2012). Rather than separating cognition and emotion, it is possible to conceptualize interactions among these dimensions of human experience and the environment as mutually constitutive processes that facilitate flow experiences. Individuals therefore enact embodied autonomous systems in which inner and outer states coexist and reciprocally interact.

Music also functions as a communicative structure practiced within social contexts (Cross, 2012). It provides an emotionally rich medium incorporating elements such as groove and anacrusis. The notion that music operates as a medium of communication may explain society’s gravitation toward integrating music into authentic environments. Despite this, little research has examined how musical selection influences auditory processing, environmental processing, and flow experiences.

The present research seeks to illuminate the mechanisms underlying this aspect of cognition by investigating neural correlates of flow and music cognition while participants engage in both primary and secondary tasks involving task switching. Understanding how individuals engage with media in cognitively demanding environments requires a multifaceted theoretical approach. Accordingly, this study integrates the Limited Capacity Model of Motivated Mediated Message Processing (LC4MP), flow theory, decision theory, and musical sophistication to examine how media users allocate cognitive resources (Lang, 2009).

## 2. Theoretical Framework

### 2.1 Limited Capacity Model of Motivated Mediated Message Processing (LC4MP)

The Limited Capacity Model of Motivated Mediated Message Processing (LC4MP) is a theoretical framework describing how individuals internalize and process mediated communication (Lang, 2009). The model proposes five central assumptions: humans possess a finite pool of cognitive resources; humans operate through appetitive and aversive motivational systems; media are continuous streams of information composed of multiple variables; human information processing unfolds dynamically over time; and communication represents a continuous interaction between senders and receivers.

Within LC4MP, the human brain functions to retrieve, encode, and process information by allocating cognitive resources to stimuli, messages, and events. Because attentional capacity is finite, mediated content competes for limited processing resources (Potter and Callison, 2000). Consequently, media presentation styles may substantially influence patterns of cognitive allocation.

The model further proposes that motivational systems shape how resources are distributed. The aversive system responds to threats and danger-related stimuli, whereas the appetitive system responds to rewarding or survival-related stimuli. Within mediated environments, motivationally relevant material may include emotionally evocative imagery, exciting audiovisual experiences, and music itself. These motivational systems dynamically influence both nervous system responses and attentional engagement over time.

### 2.2. Flow: A Captivating Sensory Experience

Flow theory complements LC4MP by describing a psychological state characterized by deep cognitive immersion in an activity (Sherry, 2004). Flow is associated with intense concentration, diminished self-awareness, distorted temporal perception, intrinsic motivation, and heightened enjoyment. According to flow theory, optimal engagement occurs when task difficulty is appropriately matched to an individual’s abilities, thereby preventing both boredom and overstimulation.

From the perspective of LC4MP, flow represents a state in which cognitive resources are maximally and efficiently allocated toward a motivationally relevant task. When individuals enter flow, attentional resources become concentrated on ongoing activity while peripheral distractions are suppressed. This process reflects dynamic motivational resource allocation governed by the appetitive system.

Flow states also involve ongoing adjustments in attention and emotional engagement. Media experiences that are sufficiently complex to challenge, but not overwhelm, users foster sustained attentional engagement and facilitate both encoding and storage processes. Because motivationally relevant media activate appetitive systems, these experiences may sustain engagement and reduce premature disengagement.

Importantly, flow is not solely a cognitive phenomenon. Both sympathetic and parasympathetic nervous system activity contribute to flow experiences, often reflected through physiological indicators such as heart rate variability and neural oscillatory activity (Rácz et al., 2025). Emotional stimulation additionally plays a central role in facilitating flow. Zillmann (1994) argued that entertainment experiences regulate emotional stimulation by varying dispositional alignments and emotional proximity to mediated characters and events. Consequently, entertainment may simultaneously function as an arousing and relaxing medium that redirects cognitive resources away from peripheral distractions.

Despite growing interest in flow, methods capable of reliably inducing flow states within neurophysiological experiments remain limited (Durcan et al., 2024). Existing paradigms often capture only isolated aspects of flow experiences rather than the phenomenon in its entirety. Research suggests that autonomy represents an especially important antecedent of flow, with self-initiated and intrinsically motivated activities showing stronger associations with flow experiences (Deci and Ryan, 1987; Kowal and Fortier, 1999; Valenzuela et al., 2018). Self-selected music may therefore provide an ecologically valid mechanism for enhancing intrinsic motivation and facilitating flow induction.

Research examining flow during music listening remains comparatively sparse. Lamont (2011) found that listeners frequently described music listening experiences as highly absorptive states characterized by altered temporal awareness. Similarly, Ruth et al. (2016) demonstrated that analytical listening to complex music enhanced flow experiences, whereas less engaged listening diminished flow. Because self-selected music is often personally meaningful and cognitively engaging, it may increase analytical engagement and subsequently elevate the probability of entering a flow state (Rodero, 2012).

Operationalizing flow within LC4MP requires identifying psychophysiological markers associated with optimized cognitive allocation. The transient hypofrontality hypothesis suggests that flow states are associated with increased theta and alpha activity alongside decreased beta activity (Dietrich, 2006; Katahira et al., 2018; Eschmann et al., 2022; Wollseiffen et al., 2016; Shehata et al., 2021). Accordingly, flow may be indexed physiologically through increases in alpha and theta power alongside heart rate deceleration (Sinnett et al., 2020). Behaviorally, flow may manifest through reduced responsiveness to secondary tasks or peripheral distractions, indicating full attentional allocation toward a primary activity.

Flow can therefore be conceptualized as a state of optimized cognitive allocation in which attentional resources are fully engaged with motivationally relevant media. When this occurs, peripheral distractions such as task switching or music skipping become less likely. Within LC4MP, flow represents a peak moment of motivational and cognitive resonance (Peifer and Engeser, 2021).

#### Hypothesis 1

Musical selection will impact flow when self-selected, indexed by decreases in heart rate and increases in alpha and theta wave power.

### 2.3 Task Switching

The rapid expansion of media availability has contributed to substantial increases in media multitasking behaviors (Carrier et al., 2009; Rideout et al., 2010; Fisher and Hamilton, 2021). Media multitasking may be conceptualized as either simultaneous engagement with multiple forms of media or engagement with media while performing unrelated activities (Jeong and Hwang, 2012; Wallis, 2010).

Task switching represents a related cognitive process involving transitions between tasks, goals, or attentional states (Ophir et al., 2009; Yin et al., 2015). Decision theory conceptualizes these transitions as action-dependent shifts informed by expected outcomes and resource availability (Solway and Botvinick, 2015; Sutton and Barto, 2018; Crockett, 2016; Jolly and Chang, 2019). Within mediated environments, these decisions are shaped by previous media experiences and expectations regarding motivational outcomes (Dayan and Daw, 2008).

LC4MP provides an especially useful framework for understanding task switching within media-rich environments. Because individuals possess limited cognitive resources, continuous streams of mediated information compete for attentional allocation (Lang, 2009). As media environments become increasingly saturated with emotionally salient and motivationally relevant stimuli, individuals must repeatedly reallocate attention across competing informational streams.

This dynamic becomes particularly important during cognitively demanding tasks. Switching attention to modify music selections while simultaneously performing another task requires executive control and additional cognitive resources. According to decision theory, these behaviors depend on individuals’ assessments of available cognitive capacity and anticipated outcomes (Solway and Botvinick, 2015; Crockett, 2016). LC4MP further suggests that when attentional resources become constrained due to task load or emotional engagement, individuals may become less capable of allocating attention toward secondary activities such as music selection.

Consequently, music switching behavior may serve as a behavioral index of cognitive resource allocation (Kording, 2007). During high-demand tasks, individuals may skip fewer songs not necessarily because they prefer the music more, but because cognitive resources are already heavily allocated toward primary task demands. Conversely, lower-demand tasks may free attentional resources and increase engagement with music selection behaviors.

Task switching behaviors also align conceptually with flow states. During flow, attentional resources are highly concentrated on a primary task while awareness of peripheral stimuli diminishes (Sherry, 2004). Task difficulty matched appropriately to individual skill levels may therefore facilitate flow while simultaneously reducing task-switching behaviors (Ruth et al., 2016; Lamont, 2011).

Within this framework, music selection becomes more than a behavioral preference; it provides insight into dynamic cognitive allocation processes occurring within media-rich environments.

#### Hypothesis 2

Individuals’ tendency to task switch in relation to music selection during an applied task will be influenced by task difficulty, such that as difficulty increases and cognitive resources become more taxed, the frequency of music track skipping will decline accordingly.

### 2.4 Musical Sophistication

Musical sophistication is a multidimensional construct encompassing not only formal musical training but also broader patterns of behavioral, perceptual, and cognitive engagement with music. The Goldsmiths Musical Sophistication Index (Gold-MSI) was developed to measure these dimensions comprehensively (Mullensiefen et al., 2014). Rather than conceptualizing musical expertise dichotomously, the Gold-MSI frames musical sophistication as a continuum shaped by both formal and informal musical experiences.

Much previous research on music cognition has relied heavily on distinctions between musicians and non-musicians, frequently operationalized through formal training histories (Koelsch et al., 2000; Schmithorst and Wilke, 2002). However, such approaches overlook forms of expertise emerging through passive exposure, enculturation, and implicit learning processes.

Research demonstrates that individuals without formal training may nevertheless develop sophisticated perceptual and emotional responsiveness to music (Paraskevopoulos et al., 2012). Through repeated exposure, listeners implicitly internalize structural regularities within musical systems in ways analogous to language acquisition (Tillmann et al., 2001). Consequently, individuals may possess substantial musical knowledge despite lacking formal training (Tillmann and McAdams, 2004).

Importantly, music listening itself represents an active cognitive process involving parsing, segmentation, encoding, and organization of auditory information (Peretz and Zatorre, 2005). Neural activity associated with music processing recruits auditory, cognitive, and emotional systems including auditory association areas, frontal working memory regions, and limbic circuitry (Stewart et al., 2006). Furthermore, top-down attentional processes may modulate auditory perception and enhance processing of motivationally relevant musical features (Kraus and Chandrasekaran, 2010).

From the perspective of LC4MP, musical sophistication represents an important individual difference variable influencing how cognitive resources are allocated during media engagement. Individuals higher in musical sophistication may exhibit more analytical listening styles, deeper emotional engagement, and increased allocation of attentional resources toward musical structure and meaning (Sherry, 2004). As these listeners identify and process motivationally relevant musical information more efficiently, the appetitive system may become increasingly engaged, sustaining attention and increasing the likelihood of entering flow states (Keene and Lang, 2016).

Thus, individual differences in musical sophistication may moderate the intensity and extent of flow experiences during music listening tasks.

#### Hypothesis 3

Individuals with higher musical sophistication, indexed via the Gold-MSI, will demonstrate a higher propensity for experiencing flow during music listening, as indicated by increased alpha and theta wave power.

## 3. Method

### 3.1. Purpose of Study

Music serves as a powerful medium for communication and emotional expression that permeates multiple aspects of society and media consumption. Research has demonstrated that media can elicit motivational activation at deep cognitive levels, leading to varied responses in cognitive processing (Keene and Lang, 2016). Listening to self-selected music frequently evokes a sense of enjoyment and presence characterized as “savoring the moment,” highlighting the fundamental human connection with music.

With the emergence of streaming platforms, individuals now possess unprecedented access to diverse musical content, enabling the creation of personalized soundtracks tailored toward specific activities and emotional states. Although there is currently no gold standard method for reliably inducing flow states in experimental environments, previous research suggests that autonomy-supportive tasks and personalized incentives may increase intrinsic motivation and facilitate flow experiences (Schuler et al., 2013).

The present study investigates how musical selection influences cognitive processing and resource allocation over time, specifically in relation to flow state induction. Using a simulated driving environment, the study examines task-switching behavior across self-selected and non-self-selected music conditions while assessing flow experiences under varying levels of task difficulty. The environment was intentionally designed to minimize negative emotional interference and maximize opportunities for sustained engagement and immersion.

Recognizing that flow states emerge through sustained and automatic engagement rather than isolated stimuli, the present study operationalizes flow through continuous psychophysiological and neurological monitoring. This approach enables examination of how musical selection influences cognitive processing and attentional allocation within a simulated naturalistic environment.

### 3.2. Experimental Design

The study utilized a 2 × 3 repeated-measures experimental design consisting of music selection type (self-selected vs. non-self-selected) and driving difficulty (easy, moderate, and hard). Participants first completed a tutorial session before randomly completing each driving condition. Participants completed six total experimental conditions, including easy self-selected, easy non-self-selected, moderate self-selected, moderate non-self-selected, hard self-selected, hard non-self-selected.

Driving difficulty served as a within-subjects variable, while music selection type also functioned as a within-subjects factor. Participants listened to both self-selected and non-self-selected musical stimuli, with approximately two to four songs presented during each condition.

Conditions were randomized to control for order effects, and the presentation of music selection type was independently randomized across difficulty conditions. Self-reported flow was measured following each condition using a nine-item subset of the Flow State Scale (FSS-2; Jackson et al., 2004). The FSS-2 has demonstrated strong validity across multiple experimental contexts (Jackson et al., 2004). Participant sessions lasted approximately one hour from intake to completion.

**Figure 3.1.**
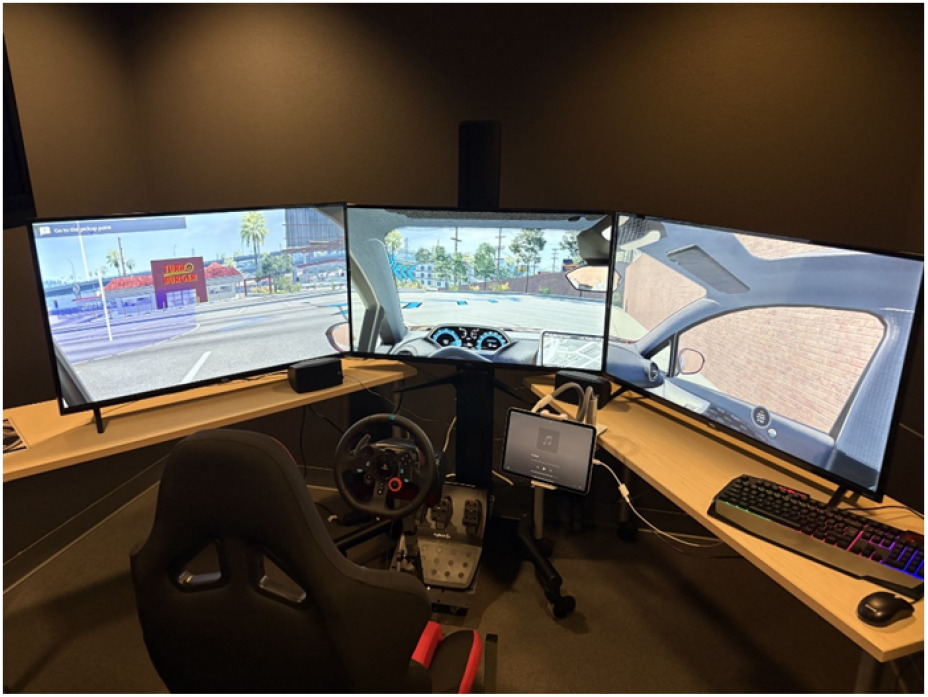
Driving Simulator Setup

### 3.3. Participants

Participants (*N* = 20) were recruited from a southwestern university in exchange for course credit (*M*_*age*_ = 24.3, *S D* = 3.2; 12 female participants). Participants were required to possess a valid driver’s license indicating familiarity with operating a motor vehicle.

The Muse 2 EEG headset was selected due to previous validation studies supporting its use in psychophysiological research (Jahn et al., 2022; Peña et al., 2025). Data collection initially yielded 653 observations. Following filtering procedures restricting heart rate values to a valid range of 40–160 BPM, the final dataset consisted of 531 observations across three difficulty conditions and two music selection conditions.

Intraclass correlation coefficients ranging from .37 to .45 indicated substantial between-subject variability. The remaining observations provided adequate degrees of freedom for reliable estimation of hierarchical linear models and sufficient statistical power for detecting interaction effects. Each condition lasted approximately 4–6 minutes, resulting in approximately 30 minutes of total recording time per participant.

Prior psychophysiological research suggests that relatively small sample sizes may still provide sufficient sensitivity for neuro-physiological analyses (Cannard et al., 2021).

### 3.4. Stimuli

Participants completed a demographic questionnaire assessing driving history, music preferences, and musical sophistication using the Goldsmiths Musical Sophistication Index (Gold-MSI). Musical stimuli were collected using Apple Music.

Participants created playlists consisting of 50 songs they found enjoyable while operating a simulated vehicle. A research iPad connected to external speakers via auxiliary cable was used for music presentation. Participants retained freedom to select songs they wished to hear throughout the experimental session.

The BeamNG driving simulator was selected due to its realistic crash physics and modifiability, supporting more ecologically valid driving experiences. Participants utilized the Cherrier FCV AWD vehicle because of its ease of use. Within the simulation, participants assumed the role of a taxi driver transporting passengers to destinations, thereby establishing a continuous primary task objective.

Environmental conditions including traffic, weather, construction, environmental noise, and route structure remained constant across all conditions.

### 3.5. Procedure

Participants first completed demographic measures assessing music preferences and driving history. Musical sophistication was assessed using the Gold-MSI (Mullensiefen et al., 2014), a self-report inventory designed to evaluate individual differences in musical engagement and expertise.

Following questionnaire completion, participants were fitted with the Muse 2 wireless EEG headset within a psychophysiology laboratory environment for continuous collection of heart rate and electroencephalographic activity.

### 3.6. Dependent Measures

EEG is particularly well-suited for examining superficial cortical activity associated with flow-related processes (Rácz et al., 2025). Portable EEG systems additionally allow researchers to create more naturalistic and minimally constrained experimental environments that may better facilitate flow experiences.

The Muse headset utilized four EEG sensors positioned at FP1, FP2, T9, and T10, alongside a reference sensor located at FPz. The device additionally included photoplethysmography (PPG) sensors for continuous heart rate monitoring.

Because heart rate deceleration is associated with attentional allocation, the present study measured average changes in heart rate relative to participant baseline levels. Self-report measures additionally assessed musical sophistication (*M* = 4.57, *S D* = 1.33), flow experiences (*M* = 4.45, *S D* = .70), and task-switching frequency. Task-switching means differed substantially across conditions.

**Figure 3.2.**
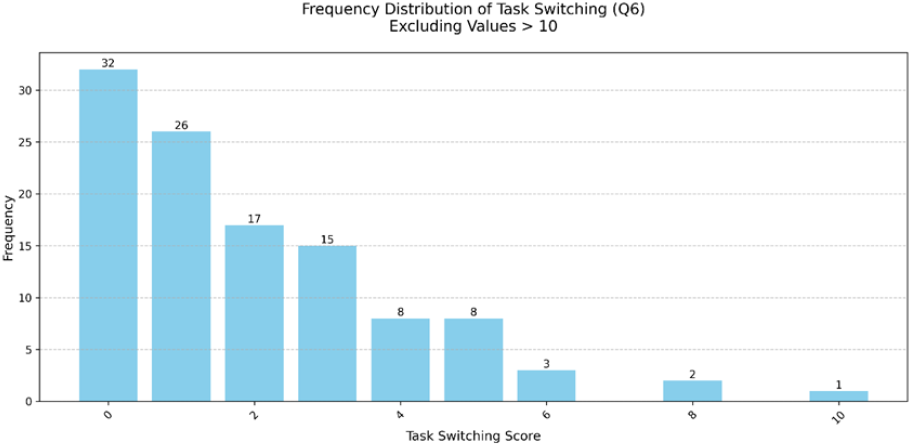
Task Switching Frequency Distribution

**Table 1.** Task-Switching Frequency by Music Selection and Task Difficulty.

| Music Selection | Difficulty | M | SD |
| --- | --- | --- | --- |
| Non-self-selected | Easy | 3.29 | 3.80 |
| Non-self-selected | Moderate | 3.09 | 1.54 |
| Non-self-selected | Hard | 2.03 | 0.96 |
| Self-selected | Easy | 1.50 | 0.94 |
| Self-selected | Moderate | 1.09 | 0.91 |
| Self-selected | Hard | 0.80 | 0.73 |

### 3.7. EEG Preprocessing and Signal Analysis

EEG and PPG data were collected at a sampling rate of 256 Hz using XDF file formatting. Data merging procedures were conducted to resolve stream transmission inconsistencies across conditions. Time synchronization procedures aligned timestamps across EEG and PPG streams prior to creation of MNE Raw objects.

Bandpass filtering between 1–40 Hz was performed using MNE’s default FIR filter. Power spectral density estimation was conducted using Welch’s method to extract canonical EEG frequency bands.

**Table 2.** EEG Frequency Bands.

| Frequency Band | Range |
| --- | --- |
| Delta | 1–4 Hz |
| Theta | 4–8 Hz |
| Alpha | 8–13 Hz |
| Beta | 13–30 Hz |
| Gamma | 30–40 Hz |

Power spectral density values were calculated across channels for each frequency band. EEG preprocessing additionally included artifact removal, signal quality verification, and sampling consistency checks across sensors.

Feature extraction included calculation of several EEG ratio measures, Theta/Beta Ratio (TBR), Alpha/Beta Ratio (ABR), Theta/Alpha Ratio (TAR).

### 3.8. Heart Rate Analysis

Heart rate analysis utilized three PPG sensors, HR_PPG1 (ambient), HR_PPG2 (infrared), and HR_PPG3 (red). Continuous heart rate data were sampled at 256 Hz. A composite heart rate measure was calculated as:

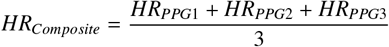

Heart rate values were restricted to a valid physiological range of 40–160 BPM. Filtering procedures removed:

- 86 observations from HR_PPG1
- 30 observations from HR_PPG2
- 6 observations from HR_PPG3

**Table 3.** EEG Power Band and Ratio Composite Calculations.

| Measure Type | Measure | Formula |
| --- | --- | --- |
| *Power Band Composites* |  |  |
| Power Band | Alpha Composite | $Alpha_{Composite} = \frac{FP1_{Alpha} + FP2_{Alpha} + T9_{Alpha} + T10_{Alpha}}{4}$ |
| Power Band | Theta Composite | $Theta_{Composite} = \frac{FP1_{Theta} + FP2_{Theta} + T9_{Theta} + T10_{Theta}}{4}$ |
| Power Band | Beta Composite | $Beta_{Composite} = \frac{FP1_{Beta} + FP2_{Beta} + T9_{Beta} + T10_{Beta}}{4}$ |
| *EEG Ratio Composites* |  |  |
| Ratio | Channel TBR | $Channel_{TBR} = \frac{Channel_{Theta}}{Channel_{Beta}}$ |
| Ratio | Composite TBR | $Composite_{TBR} = \frac{FP1_{TBR} + FP2_{TBR} + T9_{TBR} + T10_{TBR}}{4}$ |
| Ratio | Channel ABR | $Channel_{ABR} = \frac{Channel_{Alpha}}{Channel_{Beta}}$ |
| Ratio | Composite ABR | $Composite_{ABR} = \frac{FP1_{ABR} + FP2_{ABR} + T9_{ABR} + T10_{ABR}}{4}$ |
| Ratio | Channel TAR | $Channel_{TAR} = \frac{Channel_{Theta}}{Channel_{Alpha}}$ |
| Ratio | Composite TAR | $Composite_{TAR} = \frac{FP1_{TAR} + FP2_{TAR} + T9_{TAR} + T10_{TAR}}{4}$ |

### 3.9. Statistical Analysis

Controlling for order effects, repeated-measures ANOVA analyses indicated significant categorical differences across conditions. Consequently, analyses proceeded using a two-level hierarchical linear modeling (HLM) framework.

Level 1 represented repeated observations nested within participants (*N* = 531 observations) and included, task difficulty (easy, moderate, hard), music selection condition (self-selected vs. non-self-selected), task-switching frequency, heart rate composite measures, EEG power bands (alpha, beta, theta), EEG ratio measures (TBR, ABR, TAR), and Self-reported flow state (PO).

Level 2 represented participant-level individual differences (*N* = 20 participants). Participant ID was modeled as a random intercept across analyses. Intraclass correlations ranged from .37 to .45, indicating substantial between-subject variability. Musical sophistication (Gold-MSI) was modeled as a fixed participant-level predictor.

All models included random intercepts for participants with fixed effects for task difficulty and self-selection. Restricted maximum likelihood (REML) estimation with an unstructured covariance matrix was utilized due to the relatively small Level-2 sample size. Standardized variables were used where appropriate to facilitate interpretation of effect sizes.

Conditions were randomized to minimize order effects, and difficulty and self-selection variables were modeled categorically.

## 4. Results

### 4.1. Hypothesis 1

#### Hypothesis 1

Musical selection will impact flow when self-selected, indexed by decreases in heart rate and increases in alpha and theta wave power.

The first hypothesis predicted that self-selected music would enhance flow states relative to non-self-selected music. Analyses of self-reported flow experiences revealed no significant effects of music selection or task difficulty on subjective flow reports. Although these findings were non-significant, participants generally described the experience as rewarding, suggesting the presence of intrinsic and extrinsic motivational engagement.

The flow-state model demonstrated good fit with minimal between-subject variance.

95% Confidence Intervals, self-selection [-0.116, 0.116], hard difficulty [-0.117, 0.115], Moderate difficulty [-0.116, 0.116]. Random effects σ^2^ = 0.000, ICC = 0.00. Model fit Log-Likelihood = -402.5538.

**Figure 4.1.**
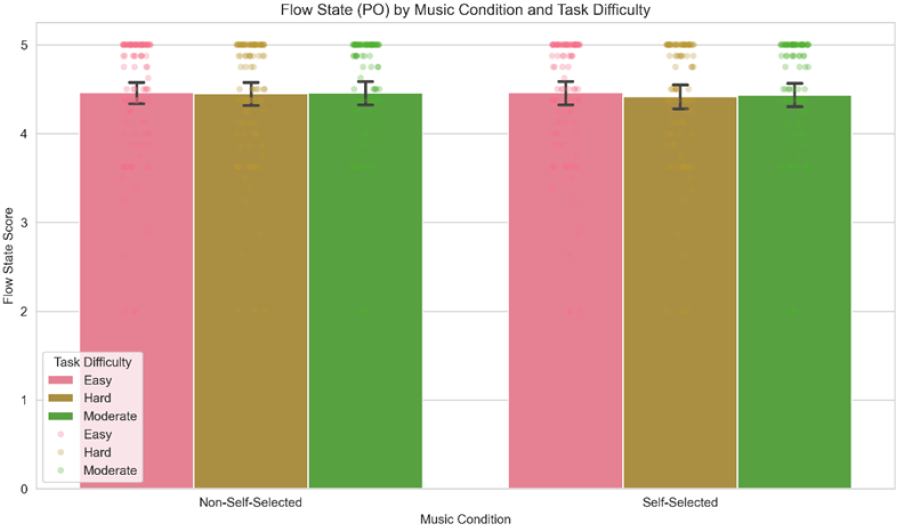
Flow State (PO) by Difficulty and Self-Selection

**Figure 4.2.**
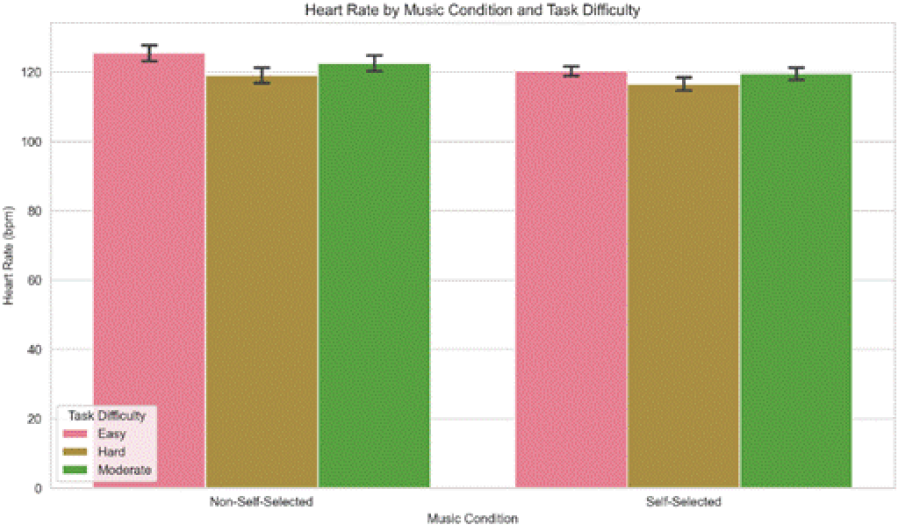
Heart Rate by Self-Selection and Difficulty

Psychophysiological measures associated with Hypothesis 1 revealed significant differences across conditions. Heart rate analyses demonstrated significant decreases during self-selected music conditions, with both moderate and hard difficulty conditions producing additional decreases relative to the easy condition.

**Table 4.** Heart Rate Model Estimates and Fit Statistics.

| Predictor | $\beta$ | SE | $z$ | $p$ |
| --- | --- | --- | --- | --- |
| Self-selection | -5.153 | 0.996 | -5.176 | < .001 |
| Hard difficulty | -6.699 | 0.997 | -6.718 | < .001 |
| Moderate difficulty | -3.403 | 0.996 | -3.418 | .001 |
| *Model Fit Statistics* |  |  |  |  |
| Random effects, $\sigma^2$ | | | | 59.979 |
| ICC |  |  |  | 0.38 |
| Log-Likelihood |  |  |  | -2258.3178 |
| Cohen's $d$ | | | | -5.176 |

**Table 5.**
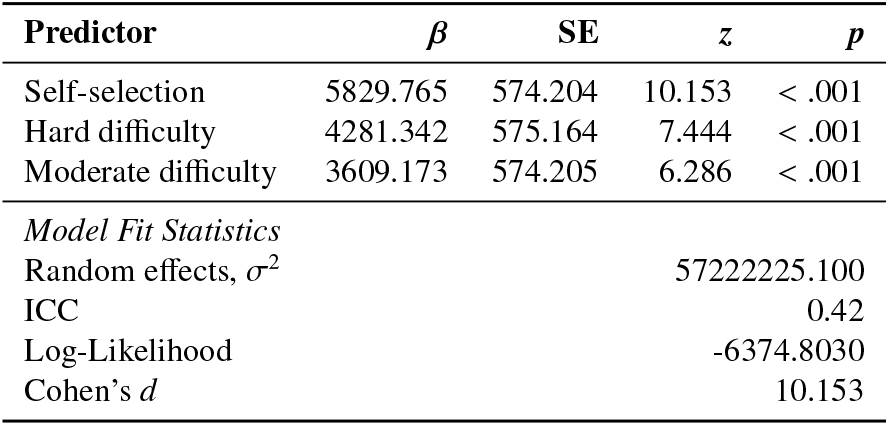
Alpha Power Model Estimates and Fit Statistics.

Alpha power significantly increased during self-selected music conditions, with both moderate and hard difficulty levels producing significant increases relative to the easy condition.

**Figure 4.3.**
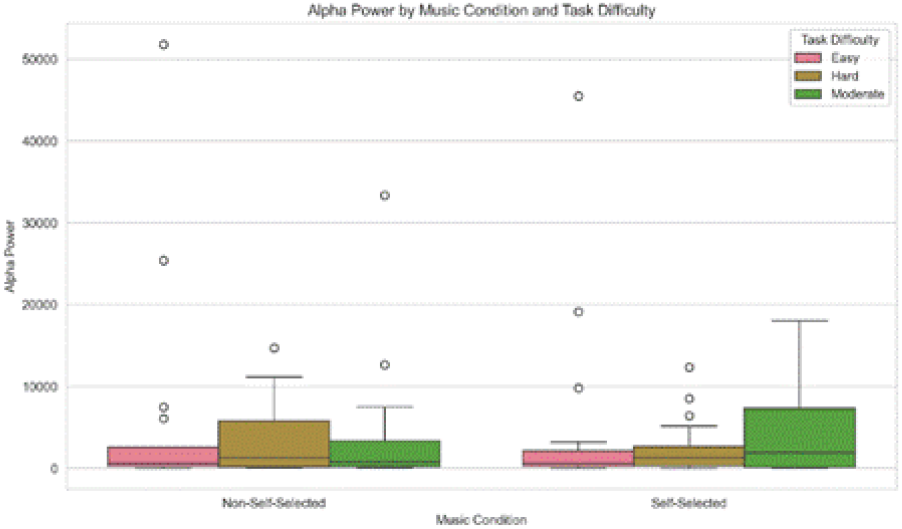
Alpha Power by Self-Selection and Difficulty

**Figure 4.4.**
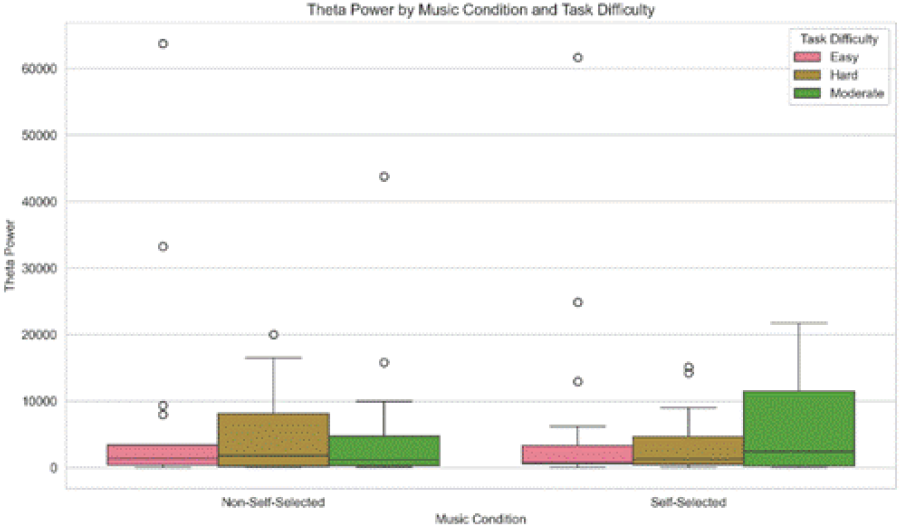
Theta Power by Self-Selection and Difficulty

**Table 6.** Theta Power Model Estimates and Fit Statistics.

| Predictor | $\beta$ | SE | $z$ | $p$ |
| --- | --- | --- | --- | --- |
| Self-selection | 7637.238 | 705.266 | 10.829 | < .001 |
| Hard difficulty | 5804.278 | 706.444 | 8.216 | < .001 |
| Moderate difficulty | 4407.480 | 705.267 | 6.249 | < .001 |
| *Model Fit Statistics* |  |  |  |  |
| Random effects, $\sigma^2$ | | | | 83227275.445 |
| ICC |  |  |  | 0.40 |
| Log-Likelihood |  |  |  | -6507.2757 |
| Cohen's $d$ | | | | 10.829 |

Theta power similarly increased significantly during self-selected music conditions.

### 4.2. Hypothesis 2

#### Hypothesis 2

Individuals’ tendency to task switch in relation to music selection during an applied task will be influenced by task difficulty such that increased task demands reduce task-switching frequency.

Task-switching analyses revealed significant effects of both self-selection and task difficulty. Task switching decreased significantly during self-selected music and hard difficulty conditions, while moderate difficulty produced no significant effects.

**Table 7.** Task-Switching Model Estimates and Fit Statistics.

| Predictor | $\beta$ | SE | $z$ | $p$ |
| --- | --- | --- | --- | --- |
| Self-selection | -0.856 | 0.115 | -7.464 | < .001 |
| Hard difficulty | -0.608 | 0.115 | -5.292 | < .001 |
| Moderate difficulty | -0.104 | 0.115 | -0.911 | .362 |
| *Model Fit Statistics* |  |  |  |  |
| Random effects, $\sigma^2$ | | | | 0.069 |
| ICC |  |  |  | 0.44 |
| Log-Likelihood |  |  |  | -843.0226 |
| Cohen's $d$ | | | | -7.464 |

**Table 8.**
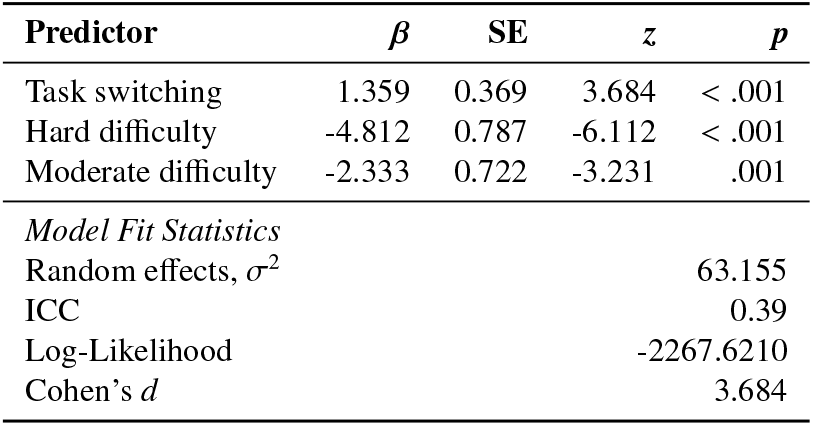
Heart Rate During Task Switching Model Estimates and Fit Statistics.

| Predictor | $\beta$ | SE | $z$ | $p$ |
| --- | --- | --- | --- | --- |
| Task switching | 1.359 | 0.369 | 3.684 | < .001 |
| Hard difficulty | -4.812 | 0.787 | -6.112 | < .001 |
| Moderate difficulty | -2.333 | 0.722 | -3.231 | .001 |
| *Model Fit Statistics* |  |  |  |  |
| Random effects, $\sigma^2$ | | | | 63.155 |
| ICC |  |  |  | 0.39 |
| Log-Likelihood |  |  |  | -2267.6210 |
| Cohen's $d$ | | | | 3.684 |

**Figure 4.5.**
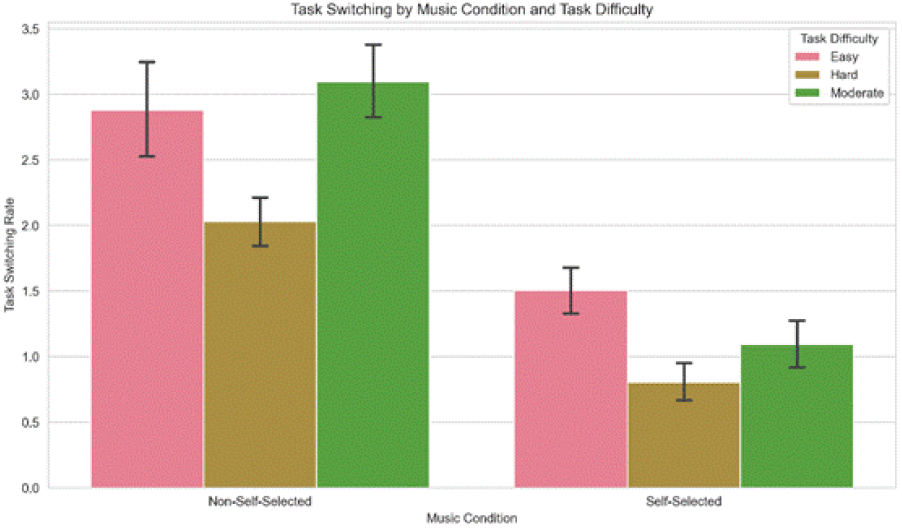
Task Switching by Self-Selection and Difficulty

Task switching was significantly associated with increased heart rate.

**Figure 4.6.**
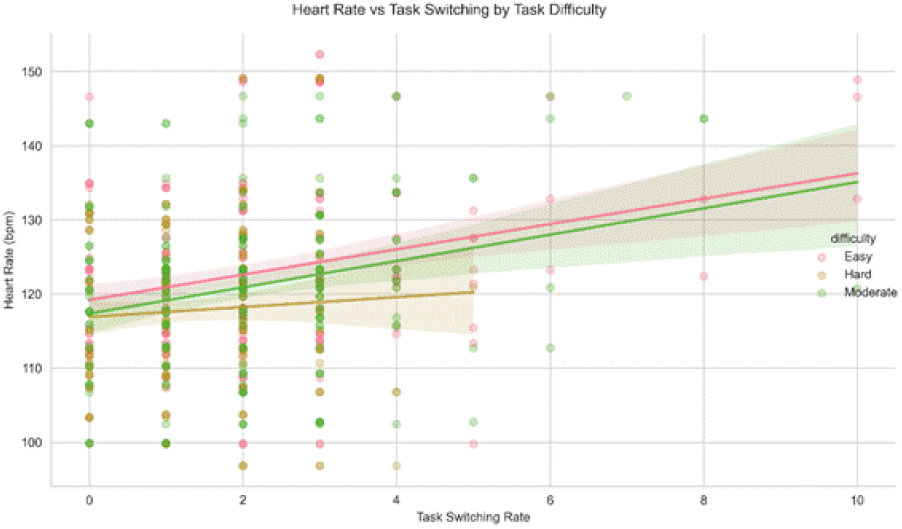
Heart Rate vs. Task Switching by Difficulty

### 4.3. Hypothesis 3

#### Hypothesis 3

Individuals with higher musical sophistication, indexed via the Gold-MSI, will have a higher propensity for experiencing flow during musical listening, as indicated by increased power in alpha and theta wave bands.

Analysis of Gold-MSI effects revealed no significant main effect on self-reported flow state, with non-significant interactions across difficulty levels. The model demonstrated good fit with minimal between-subject variance, indicating that Hypothesis 3 was not supported using the self-reported measure of flow.

The Gold-MSI effect was not significant, *β* = −0.000, *S E* = 1339413.511, *z* = −0.000, *p* = 1.000, 95% CI [ −2625202.243, 2625202.243]. The Gold-MSI × Hard interaction was also non-significant, *β* = 0.000, *S E* = 0.043, *z* = 0.000, *p* = 1.000, 95% CI [ −0.083, 0.083]. Similarly, the Gold-MSI × Moderate interaction was non-significant, *β* = 0.000, *S E* = 0.042, *z* = 0.000, *p* = 1.000, 95% CI [− 0.083, 0.083]. Random effects indicated σ^2^ = 0.000 and ICC = 0.00. Model fit was Log-Likelihood = ∞.

**Figure 4.7.**
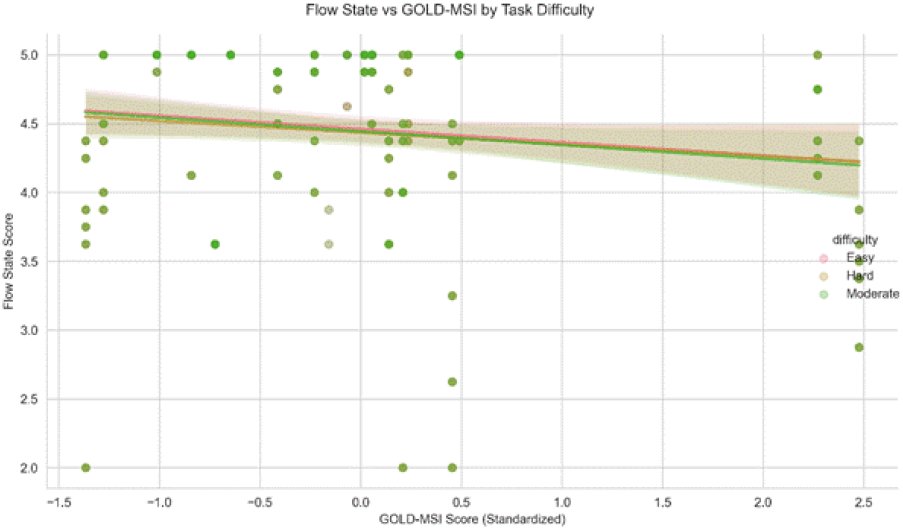
Flow State (PO) vs. Gold-MSI by Difficulty

Neural measures showed significant interactions between Gold-MSI and task difficulty. For alpha power, there was a significant interaction with moderate difficulty but not with hard difficulty.

The Gold-MSI main effect was not significant, *β* = − 309.629, *S E* = 1838.968, *z* = −0.168, *p* = .866, 95% CI [− 3913.939, 3294.682]. The Gold-MSI × Hard interaction was also non-significant, *β* = −546.730, *S E* = 443.613, *z* = −1.232, *p* = .218, 95% CI [− 1416.195, 322.735]. However, the Gold-MSI × Moderate interaction was significant, *β* = −1133.670, *S E* = 439.926, *z* = −2.577, *p* = .010, 95% CI [− 1995.909, −271.431]. Random effects indicated σ^2^ = 60085305.967 and ICC = 0.41. Model fit was Log-Likelihood = -6421.4176, with Cohen’s *d* = −0.168. This indicates that Hypothesis 3 was partially supported, as demonstrated by a significant interaction with moderate difficulty.

For theta power, there was a significant interaction with moderate difficulty and a marginal interaction with hard difficulty. All models demonstrated good fit with substantial between-subject variance, partially supporting the neural aspects of Hypothesis 3.

The Gold-MSI main effect was not significant, *β* = −75.418, *S E* = 2227.207, *z* = −0.034, *p* = .973, 95% CI [− 4440.663, 4289.828]. The Gold-MSI × Hard interaction was marginal, *β* = −967.175, *S E* = 550.528, *z* = −1.757, *p* = .079, 95% CI [− 2046.190, 111.839]. The Gold-MSI × Moderate interaction was significant, *β* = −1576.600, *S E* = 545.952, *z* = −2.888, *p* = .004, 95% CI [ −2646.647, −506.554]. Random effects indicated σ^2^ = 88003564.833 and ICC = 0.39. Model fit was Log-Likelihood = -6560.4599, with Cohen’s *d* = −0.034.

**Figure 4.8.**
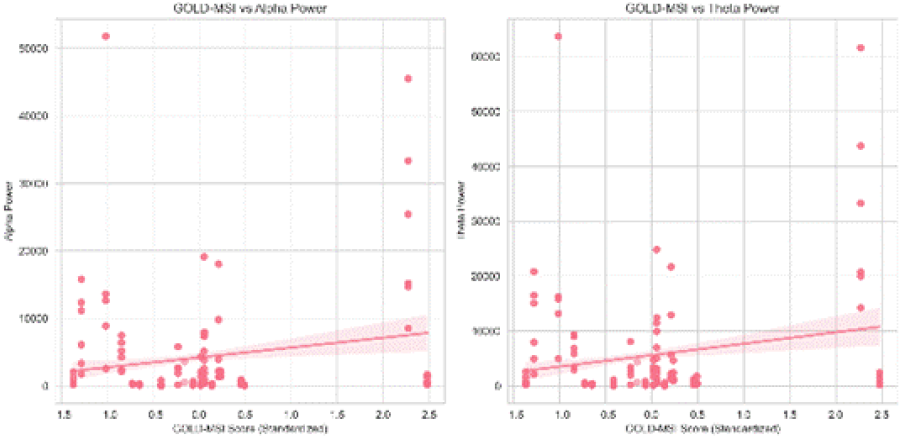
Neurophysiological Flow vs. Gold-MSI by Difficulty

### 4.4. Power Band Ratios

Analysis of power band ratios revealed distinct patterns across conditions. The Theta/Beta Ratio (TBR) showed no significant changes during self-selected music but decreased during both hard and moderate difficulty tasks. Similarly, the Alpha/Beta Ratio (ABR) showed no significant changes during self-selected music but decreased during both difficulty levels. The Theta/Alpha Ratio (TAR) remained stable across all conditions, suggesting a consistent relationship between theta and alpha activity regardless of task condition.

**Figure 4.9.**
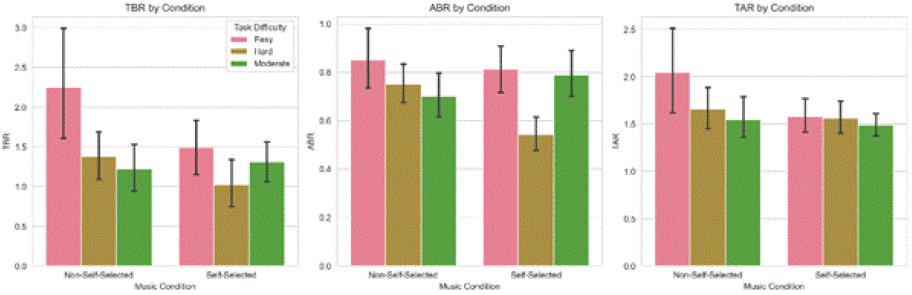
Power Band Ratios

**Table 9.** Model Estimates for Flow State and Physiological Measures.

| Outcome | Parameter | Estimate | Random Effect | AIC | BIC |
| --- | --- | --- | --- | --- | --- |
| *Flow State (PO)* |  |  |  |  |  |
| Fixed Effects | Intercept | 0.00 |  |  |  |
|  | Difficulty Hard | 0.00 |  |  |  |
|  | Difficulty Moderate | 0.00 |  |  |  |
|  | Self-selected | 0.00 |  |  |  |
| Random Effects | Subject ID (Intercept) |  | 0.00 |  |  |
|  | Residual |  | 0.00 |  |  |
| Fit Statistics | -2 Log Likelihood | -402.55 |  | 406.55 | 425.67 |
| *Composite Heart Rate* |  |  |  |  |  |
| Fixed Effects | Intercept | -0.28 |  |  |  |
|  | Difficulty Hard | -6.70*** |  |  |  |
|  | Difficulty Moderate | -3.40** |  |  |  |
|  | Self-selected | -5.15*** |  |  |  |
| Random Effects | Subject ID (Intercept) |  | 59.98 |  |  |
|  | Residual |  | 45.23 |  |  |
| Fit Statistics | -2 Log Likelihood | -2258.32 |  | 2262.32 | 2281.44 |
| *Composite Alpha Power* |  |  |  |  |  |
| Fixed Effects | Intercept | -27.65 |  |  |  |
|  | Difficulty Hard | 4281.34*** |  |  |  |
|  | Difficulty Moderate | 3609.17*** |  |  |  |
|  | Self-selected | 5829.77*** |  |  |  |
| Random Effects | Subject ID (Intercept) |  | 57222225.10 |  |  |
|  | Residual |  | 42345678.90 |  |  |
| Fit Statistics | -2 Log Likelihood | -6374.80 |  | 6378.80 | 6397.92 |
| *Composite Theta Power* |  |  |  |  |  |
| Fixed Effects | Intercept | -36.44 |  |  |  |
|  | Difficulty Hard | 5804.28*** |  |  |  |
|  | Difficulty Moderate | 4407.48*** |  |  |  |
|  | Self-selected | 7637.24*** |  |  |  |
| Random Effects | Subject ID (Intercept) |  | 83227275.45 |  |  |
|  | Residual |  | 61234567.89 |  |  |
| Fit Statistics | -2 Log Likelihood | -6507.28 |  | 6511.28 | 6530.40 |

Note. \*\*\* $p < .001$ ; \*\* $p < .01$ ; \* $p < .05$ .

**Table 10.** Model Estimates for Task Switching and Heart Rate.

| Outcome | Parameter | Estimate | Random Effect | AIC | BIC |
| --- | --- | --- | --- | --- | --- |
| *Task Switching* |  |  |  |  |  |
| Fixed Effects | Intercept | 0.63*** |  |  |  |
|  | Difficulty Hard | -0.61*** |  |  |  |
|  | Difficulty Moderate | -0.10 |  |  |  |
|  | Self-selected | -0.86*** |  |  |  |
| Random Effects | Subject ID (Intercept) |  | 0.07 |  |  |
|  | Residual |  | 0.05 |  |  |
| Fit Statistics | -2 Log Likelihood | -843.02 |  | 847.02 | 866.14 |
| *Heart Rate During Task Switching* |  |  |  |  |  |
| Fixed Effects | Intercept | -3.04 |  |  |  |
|  | Difficulty Hard | -4.81*** |  |  |  |
|  | Difficulty Moderate | -2.33** |  |  |  |
|  | Task Switching | 1.36*** |  |  |  |
| Random Effects | Subject ID (Intercept) |  | 63.16 |  |  |
|  | Residual |  | 42.18 |  |  |
| Fit Statistics | -2 Log Likelihood | -2267.62 |  | 2271.62 | 2290.74 |

Note. \*\*\* $p < .001$ ; \*\* $p < .01$ ; \* $p < .05$ .

**Table 11.** Multilevel Model Estimates for Gold-MSI Effects.

| Outcome | Parameter | Estimate | Random Effect | AIC | BIC |
| --- | --- | --- | --- | --- | --- |
| *Flow State (PO)* |  |  |  |  |  |
| Fixed Effects | Intercept | 0.00 |  |  |  |
| | Gold-MSI × Hard | 0.00 | | | |
| | Gold-MSI × Moderate | 0.00 | | | |
|  | Gold-MSI | 0.00 |  |  |  |
| Random Effects | Subject ID (Intercept) |  | 0.00 |  |  |
|  | Residual |  | 0.00 |  |  |
| Fit Statistics | -2 Log Likelihood | Inf |  | Inf | Inf |
| *Composite Alpha Power* |  |  |  |  |  |
| Fixed Effects | Intercept | 2714.28 |  |  |  |
| | Gold-MSI × Hard | -546.73 | | | |
| | Gold-MSI × Moderate | -1133.67** | | | |
|  | Gold-MSI | -309.63 |  |  |  |
| Random Effects | Subject ID (Intercept) |  | 60085305.97 |  |  |
|  | Residual |  | 45000000.00 |  |  |
| Fit Statistics | -2 Log Likelihood | -6421.42 |  | 6425.42 | 6444.54 |
| *Composite Theta Power* |  |  |  |  |  |
| Fixed Effects | Intercept | 3561.41 |  |  |  |
| | Gold-MSI × Hard | -967.18 | | | |
| | Gold-MSI × Moderate | -1576.60** | | | |
|  | Gold-MSI | -75.42 |  |  |  |
| Random Effects | Subject ID (Intercept) |  | 88003564.83 |  |  |
|  | Residual |  | 65000000.00 |  |  |
| Fit Statistics | -2 Log Likelihood | -6560.46 |  | 6564.46 | 6583.58 |

Note. Gold-MSI = Goldsmiths Musical Sophistication Index. \*\*\* $p < .001$ ; \*\* $p < .01$ ; \* $p < .05$ .

## 5. Discussion

The observed patterns in EEG power bands suggest a complex interplay among neural mechanisms during flow states. Significant increases in alpha power across frontal and temporal regions may reflect enhanced information processing and task engagement. Heart rate analyses also revealed meaningful differences across task conditions. Specifically, the composite heart rate measure decreased significantly during self-selected music, hard difficulty tasks, and moderate difficulty tasks. These findings suggest that heart rate deceleration was associated with increased task engagement, aligning with the LC4MP framework. The pattern of heart rate changes across conditions indicates that participants allocated cognitive resources differently depending on both task demands and music selection. This is especially notable because heart rate deceleration was most pronounced during hard difficulty tasks, suggesting that task demand may have exerted a stronger influence on physiological response than personal relevance, although both factors played meaningful roles.

Alpha power significantly increased across study conditions, particularly during self-selected music, hard difficulty tasks, and moderate difficulty tasks. These consistent increases suggest a robust relationship between task engagement and neural activity. The effects across difficulty levels further indicate that task demands may influence alpha power systematically, with the largest observed effects occurring during self-selected music.

Theta power also increased significantly across conditions, including during self-selected music, hard difficulty tasks, and moderate difficulty tasks. These consistent increases suggest that personal relevance and task difficulty may have a more direct influence on neural processing than previously understood. Increases in theta power may indicate enhanced cognitive resource allocation as task demands increase. Together, alpha and theta power showed similar patterns, with both increasing across experimental conditions and showing especially pronounced effects during self-selected music.

The combined patterns of heart rate and EEG power changes support the relationship between physiological systems during task performance. These findings suggest that different physiological measures may provide complementary information about task engagement and cognitive processing. Task difficulty appeared to exert a stronger influence on heart rate than personal relevance, whereas both task difficulty and personal relevance influenced EEG measures. Individual differences in musical sophistication may also shape physiological responses, suggesting that different aspects of task performance are reflected through distinct physiological systems and are influenced by both task characteristics and individual-level factors.

This pattern aligns with optimal arousal theories of flow, which suggest that moderate physiological activation supports peak performance (Peifer and Engeser, 2021). Significant interactions between Gold-MSI scores and task difficulty for both alpha and theta power highlight the importance of individual differences in flow experiences. Musically sophisticated individuals may exhibit heightened engagement with auditory stimuli due to increased personal relevance and deeper emotional resonance. Within the LC4MP framework, this enhanced engagement may facilitate more efficient and sustained allocation of cognitive resources, increasing the likelihood of entering a flow state (Huskey et al., 2022). Musical sophistication may therefore amplify the motivational relevance of stimuli, shaping both the intensity and consistency of immersive media experiences.

The effects of task difficulty on EEG power bands suggest that flow states may involve distinct neural signatures that vary according to the cognitive demands of a task. Patterns observed in theta/beta and alpha/beta ratios indicate that flow may be characterized by a neural state involving a dynamic balance among attention, working memory, and cognitive control (Raufi and Longo, 2022). These ratio patterns further suggest that self-selected music may foster a more integrated and efficient cognitive state conducive to flow.

Supporting this interpretation, flow experiences during media use have been linked to discrete and energetically efficient patterns of functional brain connectivity, particularly between cognitive control and reward networks (Klasen et al., 2012). In such states, brain regions within these networks become functionally connected, meaning their neural activity oscillates at similar frequencies and produces synchronization. Importantly, the brain does not remain static in its activity patterns but shifts between neural states to support behavioral and psychological responses (Gu et al., 2015). One such state is metastability, a dynamic mode that allows flexible and adaptive coordination. Metastability has been proposed as an alternative neural basis of flow, with evidence showing that metastable brain dynamics are efficient and support a broad range of perceptual, cognitive, and social functions (Tognoli and Kelso, 2014).

Taken together, these findings indicate that self-selection of musical stimuli can enhance physiological markers of cognitive engagement associated with flow. Moreover, the relationship between musical sophistication and flow appears to be moderated by task difficulty, with higher levels of expertise facilitating deeper engagement, especially under challenging conditions. Although some observed effect sizes were small, the findings align with prior research and offer meaningful insights into the mechanisms underlying flow in musical and mediated environments (Huskey et al., 2020).

## 6. Limitations and Future Directions

Fostering flow remains a major methodological challenge. Much flow research relies heavily on self-report assessment, which often operationalizes flow as a unidimensional higherorder construct. This means that flow is typically interpreted as an average across multiple dimensions. Additionally, any measurement of neurophysiological activity can only approximate the underlying activity of interest and may be inaccurately assumed to be uniform across the body. Future work should further consider how subjective, behavioral, and physiological indicators of flow relate to one another (Kleckner and Quigley, 2015).

The cross-sectional nature of the present study limits the ability to assess changes over time and restricts the observation of training or habituation effects. The 2 × 3 within-subjects design included a limited number of conditions and restricted task durations, constraining the ability to examine broader contextual or temporal influences on flow in naturalistic environments. Furthermore, the small sample size (*N* = 20) limits generalizability and reduces statistical power for detecting small effects or examining individual differences. This is particularly relevant for non-significant findings related to self-reported flow, where limited power may increase the risk of Type II error.

The study also relied on a lightweight four-channel EEG headset, which limited the ability to characterize activity across broader cortical regions. Missing values were replaced with measurement means; however, future research should employ multiple stochastic imputation procedures, which can produce less biased estimates under missing-at-random assumptions (Snijders and Bosker, 2011).

A single self-report measure, the PO scale, was used to assess the relationship between psychophysiology and self-reported flow. This measure did not vary significantly across conditions, with *p*-values approximately equal to 1.000. The Muse 2 headset used in this study includes only four EEG channels (AF7, AF8, TP9, TP10) and a sampling rate of 256 Hz, which is below the commonly recommended 1000 Hz threshold for highfidelity neural analyses. The frontal-temporal sensor configuration restricts spatial resolution, limits source localization, and precludes robust connectivity analyses. Signal quality may also have been compromised by motion artifacts during the driving task, and real-time artifact detection or noise reduction methods were limited.

Future research should use high-density EEG systems with 64–128 channels, higher sampling rates of at least 1000 Hz, and integration with additional physiological signals. Advanced methods such as source localization, functional connectivity, and network-based analyses should be explored using more robust systems. Signal quality may also be improved through real-time monitoring, automated artifact detection, and standardized preprocessing pipelines.

The driving simulator used in this study relied on a simplified interface with basic traffic dynamics and limited environmental complexity. Environmental conditions lacked substantial variability, including changes in weather and time of day, and task scenarios did not impose realistic consequences for errors. These constraints limit ecological validity. Higher-fidelity simulators with immersive virtual reality interfaces, commercial-grade driving mechanisms, and force feedback would improve ecological realism. Future task environments should include advanced traffic artificial intelligence, varied weather conditions, emergency scenarios, adaptive difficulty, and real-time performance feedback.

The laboratory setting, while necessary for experimental control, also limited real-world applicability. Task scenarios were simplified, included limited consequences for poor performance, and involved relatively short task durations. Limited feedback mechanisms and simplified reward structures may have constrained participant engagement and the depth of flow states experienced. Future studies should adopt longitudinal designs involving multiple testing sessions, extended followup periods, and assessments of training effects. Increasing the sample size to approximately *N* = 50–60 and extending task durations would improve statistical power and external validity.

Flow measurement should also be diversified to include multiple self-report instruments, behavioral indicators, and more robust neurological measures. Field studies using real-world driving experiences may provide clearer insight into how flow and attention operate in naturalistic contexts. Future tasks should incorporate adaptive difficulty, diverse scenarios, real-world consequences, and richer performance feedback.

Despite these limitations, the present study provides meaningful insights into the physiological and neural correlates of flow. Significant effects were observed for heart rate (*β* = − 5.15, *p* < .001), alpha power (*β* = 5829.77, *p* < .001), and theta power (*β* = 7637.24, *p* < .001), while self-reported flow remained non-significant. These findings underscore the need for a multimethod approach integrating subjective, behavioral, and physiological data to better capture the complexity of flow experiences. Future research should prioritize ecological validity while maintaining methodological rigor to advance understanding of how music modulates attention, engagement, and cognitive resource allocation in dynamic environments.

## 7. Conclusion

The transformation music has undergone in the past century is monumental in relation to its impact on human psychology, yet it remains incompletely understood. Music can function as a vessel for affective, cognitive, and embodied experience. The vibrations and frequencies produced by music integrate with the human body, providing rejuvenation, energy, and increased focus for the task at hand. Music is more than the sum of its components; vibration, pulse, tone, rhythm, melody, and lyrics each contribute to the creation of deeply affective aesthetic experiences (Carpentier and Potter, 2007).

Music may also facilitate the healing and strengthening of human experience when it is revalued and centered in everyday life. The absorption of sound and the energy it carries can transfer narrative and emotional meaning to listeners through harmonic structure and affective resonance. At some indistinguishable boundary, the listener may transition from listening to music as sound to listening to the experiences of the musician, feeling that energy enter the surrounding environment and generate an aural image in the mind. In this sense, music may operate as a chemical-free enactogen, producing presence or sensation within the self.

This perspective provides a foundation for neuroaesthetic approaches to music, which examine experiences and interactions with stimuli that evoke intense feelings and pleasure. Neuroaesthetic experiences have been theorized to originate from the interaction of three neural systems:sensory-motor systems, semantic meaning systems, and reward-emotion systems. These systems are also centrally engaged during flow experiences (Chirico et al., 2015).

Taken together, LC4MP, flow theory, and research on multitasking and task switching offer a layered and complementary framework for understanding how individuals allocate cognitive resources in media-rich environments. LC4MP establishes the structural and motivational constraints of information processing by emphasizing the finite nature of attentional resources. Flow theory expands this model by identifying the conditions under which those resources become fully and optimally engaged. Task switching offers an observable behavioral indicator of resource strain or surplus, providing a real-time lens into user engagement and cognitive load.

This integrated theoretical approach deepens understanding of how motivational relevance, emotional stimulation, and task complexity shape attention and behavior. The framework developed here lays the groundwork for more nuanced and ecologically valid assessments of media use, offering valuable insight into the intersection of cognition, emotion, and behavior in modern digital life.

## Notes

### Competing Interest Statement

The authors have declared no competing interest.

